# A chromosome-level assembly of the *Fusarium oxysporum* biocontrol strain FO12

**DOI:** 10.64898/2026.03.24.713715

**Authors:** Andrea Doddi, Ana López-Moral, Hayley Mangelson, Antonio Di Pietro, Carlos Agustí-Brisach

## Abstract

*Fusarium oxysporum* FO12 was originally isolated from cork oak (*Quercus suber* L.) and has been characterised as a highly effective biological control agent of wilt diseases on different crops. FO12 endophytically colonises roots and basal stems of plants, reducing the establishment of the soil-borne pathogen *Verticillium dahliae* and triggering plant defence-related genes. Here, we report a chromosome-level genome assembly of FO12 using Nanopore and Hi-C data. The 57.60 Mb assembly comprises 14 chromosome-scale scaffolds with centromeres resolved and telomeric repeats detected at 4 of 28 chromosome ends. This high-quality reference genome provides a valuable resource for further research into the use of FO12 in agriculture as a biocontrol agent.

## Background & Summary

The filamentous fungus *Fusarium oxysporum* species complex includes a large number of lineages, that have been classified into different formae speciales (ff. spp.) based on their capacity to cause vascular wilt on more than 100 crop species in a host-specific manner^1–3^. Strains of the *Fusarium oxysporum* species complex isolates are also natural producers of secondary metabolites and toxins such as fusaric acid, posing threats to plant and human health^4^. Although notorious as plant pathogens, *F. oxysporum* strains are also found as plant endophytes, often with beneficial effects for the host plant^5,6^. The genome of *F. oxysporum* is compartmentalised into conserved core chromosomes and lineage-specific (LS) accessory chromosomes or regions^7^. Accessory regions are enriched in transposable elements (TEs) and display a low gene density and a high GC content compared to core regions^7^, making sequence assembly highly challenging. Accessory chromosomes have been associated with host-specific virulence and niche adaptation but are also found in non-pathogenic endophytic strains^7–10^.

Recent population genomic studies revealed significant structural variation and evolutionary dynamics in the accessory compartments of the *F. oxysporum* species complex^10^. These findings indicate that 11 core chromosomes are consistently shared among all *F. oxysporum* strains, while the number and genomic content of the accessory chromosomes differ between isolates but tend to be similar within the same forma specialis^7–10^.

*F. oxysporum* strain FO12 was originally isolated from cork oak (*Quercus suber* L.) and has been reported as a beneficial endophyte acting as a highly effective biological control agent and biofertilizer on different woody (e.g. olive) and herbaceous crops (e.g. cotton, cucumber, tomato)^11–14^. Previous studies demonstrated that FO12 can endophytically colonise the roots and basal stems of woody and herbaceous plants, reducing or preventing the establishment of the soil-borne wilt pathogen *Verticillium dahliae*^11,12^. FO12 was also shown to trigger expression of plant genes involved in Induced Systemic Resistance (ISR), Systemic Acquired Resistance (SAR), and iron deficiency response^13^. Collectively, these studies suggest a lack of pathogenic behaviour of this strain. However, the molecular mechanisms underpinning its beneficial effect and the absence of pathogenicity remain poorly understood.

Here we report the first chromosome-level genome assembly of FO12, based on a hybrid sequencing approach. Long-read Oxford Nanopore sequencing generated 7.03 Gb of high-quality reads, providing approximately 123× raw genome coverage. Reads ≥ 3 kb were selected to achieve an optimal effective coverage for the initial *de novo* assembly, which was performed using Flye v. 2.9.6^15^. To achieve chromosome-level continuity, Hi-C scaffolding was incorporated. The Hi-C library generated 34.31 Gb of raw data via 150-bp paired-end (PE150) sequencing (114,372,691 read pairs), providing an estimated 595× genome coverage. The data were processed via Juicer v2.0^16^ to obtain 66,927,414 valid, unique chromatin interaction pairs (Table 1). Scaffolding was subsequently executed using 3D-DNA^17^ to construct the final pseudomolecules. As a final step, manual curation was carried out using Juicebox^18^.

**Table 1.** Assembly statistics and sequencing data of the *Fusarium oxysporum* strain FO12 genome.

| Assembly Features |  |
| --- | --- |
| Total assembly length (bp) | 57605083 |
| Total number of scaffolds | 46 |
| Resolved chromosomes (Core / Accessory) | 14 (10 / 4) |
| Largest scaffold (Mb) | 11.15 |
| Scaffold N50 (Mb) | 4.77 |
| Scaffold L50 | 5 |
| Overall GC content (%) | 47.63 |
| Sequencing Data |  |
| Oxford Nanopore reads (Gb) | 7.03 (~123xcoverage) |
| Hi-C Illumina PE150 data (Gb) | 34.31 (~595x coverage) |
| Hi-C sequenced read pairs | 114372691 |
| Hi-C valid chromatin interactions | 66927414 |

The hybrid assembly pipeline produced 46 scaffolds totalling 57,605,083 bp and 14 chromosome-scale scaffolds, as supported by high-depth Hi-C interaction blocks. The use of Hi-C centromeric interaction blocks is standard in fungal genome analysis to define chromosome boundaries in cases where telomere-to-telomere assembly is not achieved^19^. Since only 4 of the 28 telomeres were detected (TRF/StainedGlass), we report here chromosome-scale scaffolds rather than telomere-to-telomere assemblies. The assembly exhibits continuity metrics, with a scaffold N50 of 4.77 Mb, an L50 of 5, and the largest scaffold reaching 11.15 Mb (Figure 1A; Table 1; Supplementary Table 1). The overall genome architecture visualised by circos plot (Fig. 1B) highlights the heterogeneous distribution of genomic features. In line with the compartmentalised genome structure of *F. oxysporum*, the conserved core chromosomes shared across the *F. oxysporum* species complex^7,10^ are clearly conserved.

**Figure 1.**
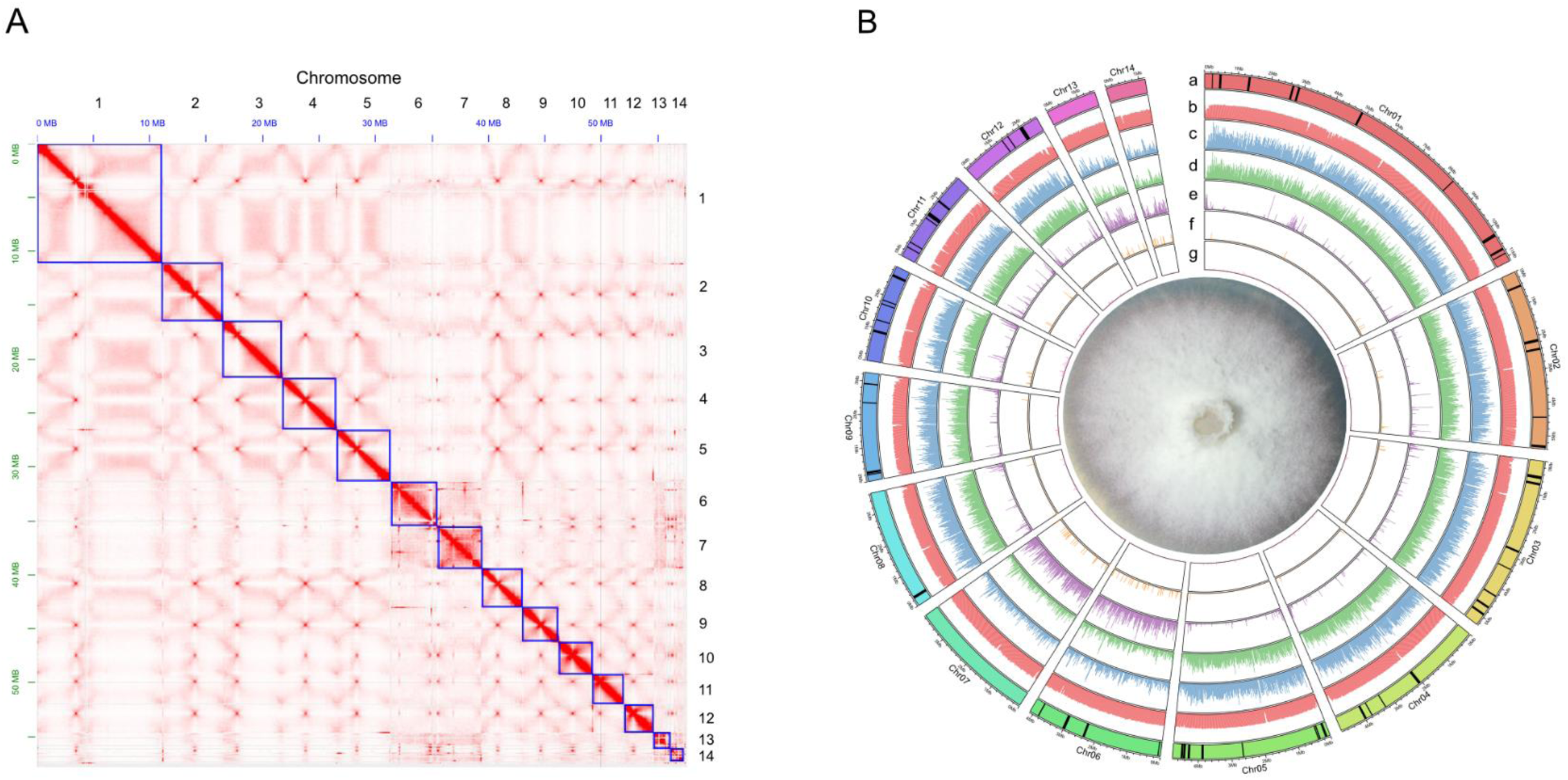
Overview of the chromosome-level genome assembly and annotation of *Fusarium oxysporum* strain FO12. **(A)** Genome-wide Hi-C contact map showing the interaction matrix among the 14 assembled chromosomes. **(B)** Circular plot illustrating the genomic features of the FO12 genome. Tracks from outer to inner represent: (a) Ideograms of the 14 chromosomes, in which the black vertical lines represent the genomic locations of biosynthetic gene clusters (BGCs); (b) GC content; (c) gene density; (d) exon density; (e) repetitive DNA (transposable elements) density; (f) simple repeat density; and (g) ncRNA (non-coding RNA) density. Genomic feature densities (tracks b to g) were calculated in non-overlapping 20 kb windows and represented on a scale from 0 to 1, except for track g, which ranges from 0 to 0.1. Chr: chromosome

In addition, FO12 has four accessory chromosomes which exhibit significantly lower gene density and a substantial enrichment in TEs. Moreover, biosynthetic gene clusters (BGCs) and non-coding RNAs are arranged in specific spatial configurations, often located near TE-rich boundaries, thereby reflecting the bipartite genome architecture typical of the *F. oxysporum* species complex (Fig. 1B).

To further characterise the chromosome set of FO12 we performed whole-genome sequence alignments with the tomato pathogenic isolate Fol4287 (GCA_000149955.2) and the non-pathogenic isolate Fo47 (GCA_013085055.1), using minimap2^20^. This revealed that FO12 shares the core chromosome regions with Fo47 and Fol4287, although we noted that Chr01 of FO12 is significantly larger than the homologs of Fo47 and Fol4287. The remaining region of FO12 Chr01 aligns almost completely with core chromosomes Chr08 and Chr06 of Fol4287 and Fo47, respectively, suggesting the occurrence of a core chromosome fusion event in FO12 (Figure 2A).

**Figure 2.**
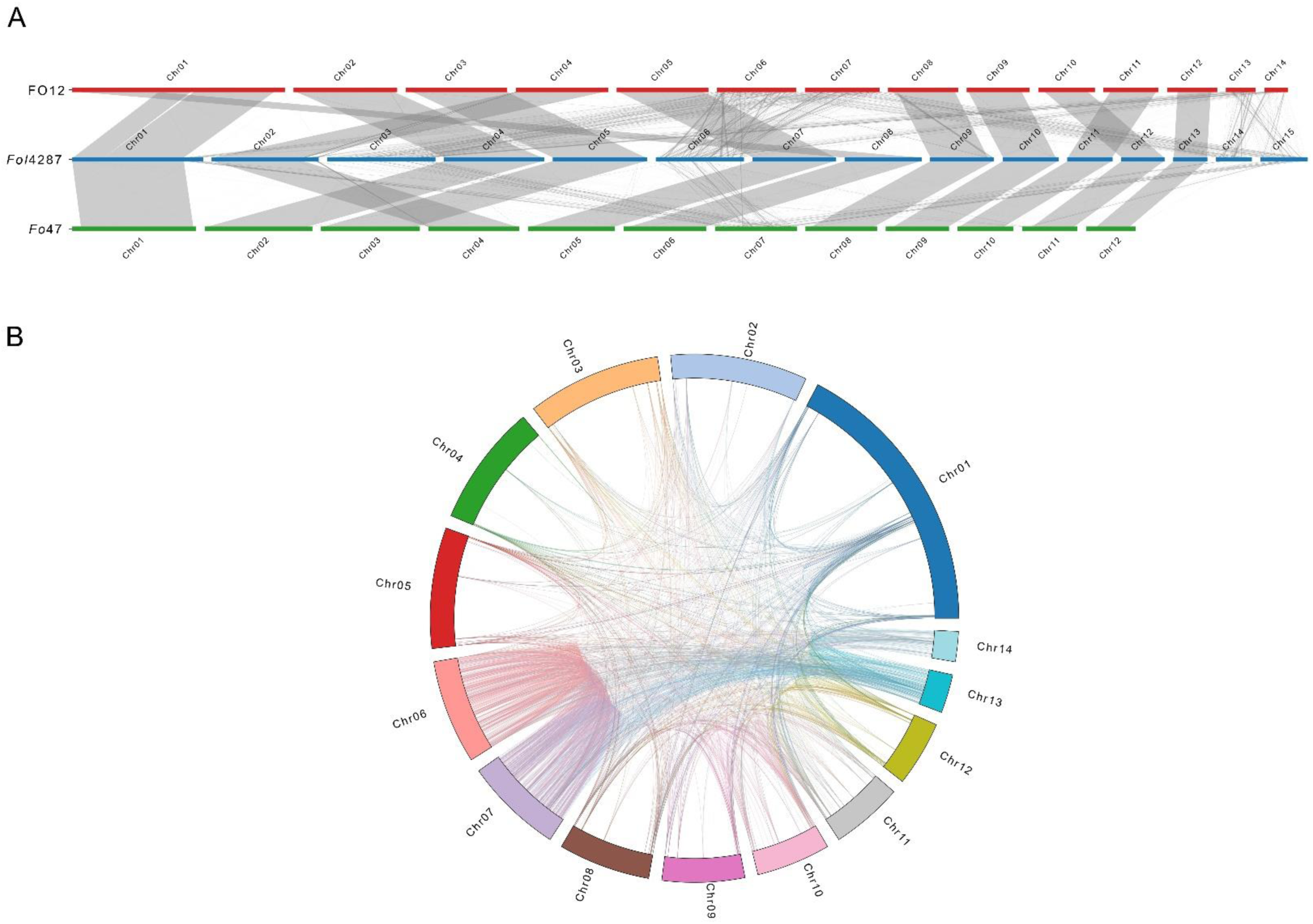
The genome of *Fusarium oxysporum* FO12 exhibits evidence for chromosome fusion and duplication events. **(A)** Linear synteny plot comparing the genomic architectures of the isolate FO12 (top, red), the tomato pathogen reference strain Fol4287 (middle, blue), and the non-pathogenic biocontrol strain Fo47 (bottom, green). Horizontal bars represent individual chromosomes, while grey ribbons represent syntenic blocks. **(B)** The circos plot illustrates intra-genomic duplications within the FO12 genome. The outer ring displays the FO12 chromosomes, while the internal colored links represent extensive self-alignments and segmental duplications.

The four accessory chromosomes of FO12, Chr06, 07, 13 and 14, encompass a total of 10.70 Mb, corresponding to 18.6% of the total genome size (Figure 2A). The fraction of accessory genome in FO12 is larger than in Fo47, which only contains a single accessory chromosome (Chr06) of 4.25 Mb, but smaller than in Fol4287, whose accessory chromosomes share only a low degree of similarity with the accessory Chrs 03, 14 and 15 of FO12 (Figure 2A).

Accessory Chr07 and Chr06 of FO12 share an average identity of 89.0%, suggesting that may have originated from a duplication event (Supplementary Table 2; Supplementary Image 1). Accessory Chr13 and Chr14 could represent secondary duplication events of Chr06 and Chr07, followed by partial loss of the duplicated regions (Figure 2B; Supplementary Table 2; Supplementary Image 1), although horizontal transfer or assembly artefacts cannot be ruled out.

Ab initio and homology-only evidence predicted 16,068 protein-coding genes and 319 tRNAs. Transcript-guided refinement may further adjust this number. The protein-coding genes span a total of 24,122,300 bp with an average gene length of 1,472 bp (Table 2). Functional annotation was performed using a multi-tiered approach against several public databases, including UniProtKB/Swiss-Prot, KEGG, InterPro, Pfam, and Gene Ontology (GO). Conserved protein domains, functional motifs, and biological pathways (GO, KEGG, Pfam, and InterPro) were primarily annotated through the Funannotate pipeline v1.8.1^21^, which integrates the predictive power of InterProScan^22^ and eggNOG-mapper^23^. To robustly validate the predicted functions, the curated protein sequences were aligned against the manually annotated Swiss-Prot database using Diamond v2.1.10 (parameters: --evalue 1e-5, --sensitive). Overall, this strategy successfully assigned functional information to 97.5% (15667) of the predicted proteins across at least one public database (Figure 3, Table 4).

**Figure 3.**
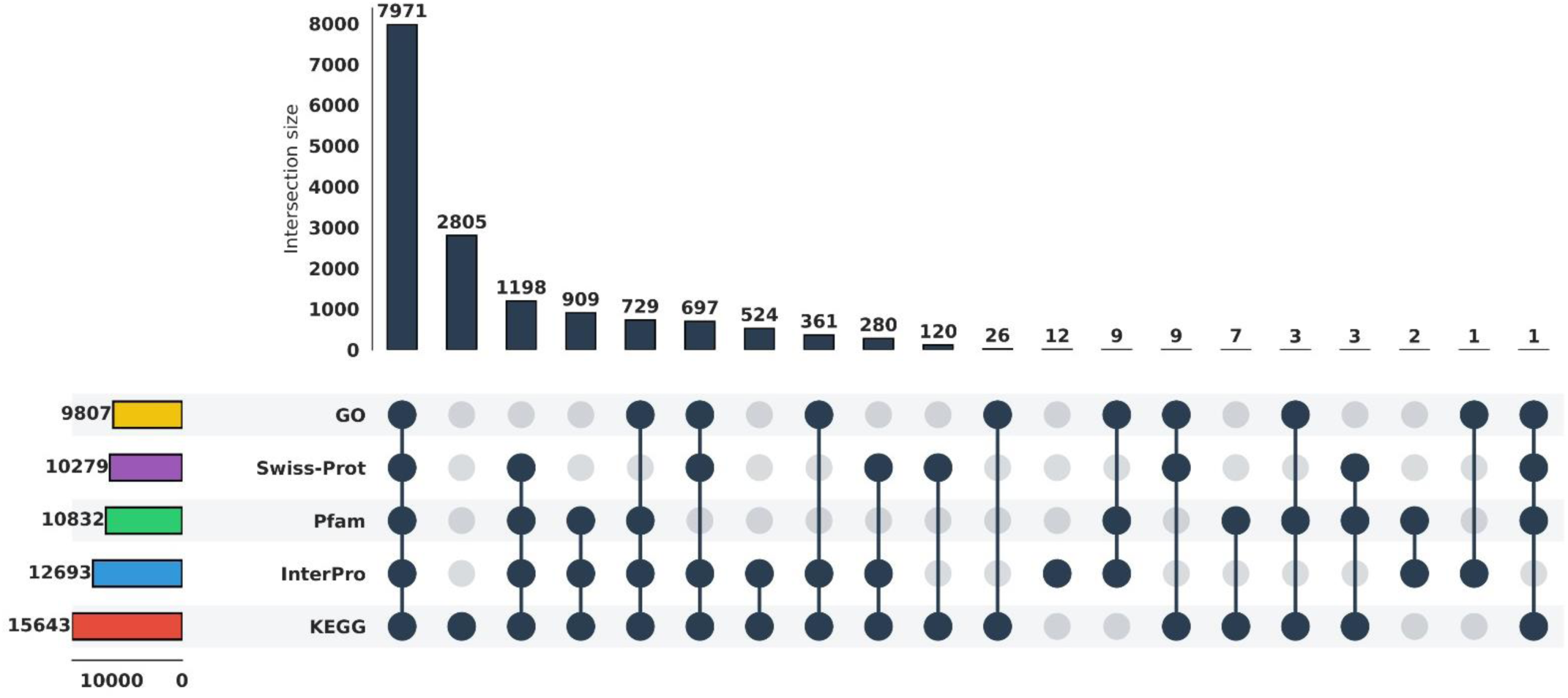
Functional annotation of predicted genes in the genome of *Fusarium oxysporum* strain FO12. The upset bar plot visualises the functional annotation results based on different databases.

**Table 2.**
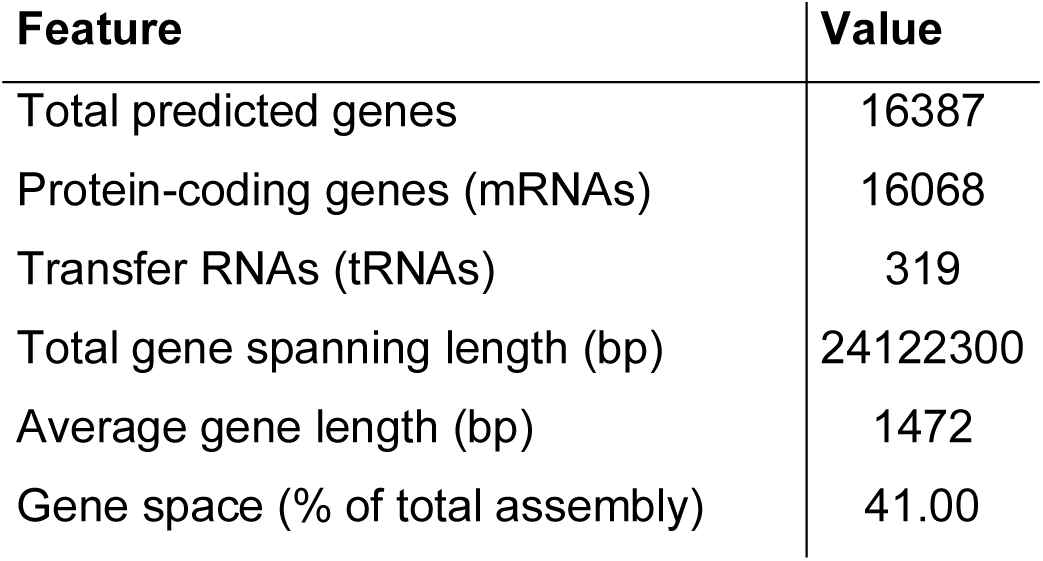
Coding gene prediction of *Fusarium oxysporum* strain FO12.

TEs were previously shown to act as major drivers of adaptive evolution in *F. oxysporum*^24^. A *de novo* TE annotation, including a curated TE library from Fol4287^7^, revealed that TEs occupy 5.75 Mb, corresponding to 9.98% of the FO12 genome (Table 3; Supplementary Table 3). Among the classified elements, Long Interspersed Nuclear Elements (LINEs) are the most abundant class, comprising 1.23% of the total genome, with the Tad1 family being the dominant subgroup with 1.17%. DNA transposons of the DNA/DTM class show a similarly high prevalence with 1.14%, followed by Long Terminal Repeat (LTR)/Copia retrotransposons (0.81%) and DNA/DTA retrotransposons (0.60%). LTR/Gypsy retrotransposons, including characteristic elements such as Skippy, make up a smaller fraction (0.11%) of the genome (Table 3).

**Table 3.** Statistics of repeat sequences in the *Fusarium oxysporum* strain FO12 genome.

| TE Class | Count | Total Masked(bp) | Percent Genome(%) |
| --- | --- | --- | --- |
| Unspecified | 3236 | 2694446 | 4.68 |
| LINE/Tad1 | 561 | 675025 | 1.17 |
| DNA/DTM | 955 | 654592 | 1.14 |
| LTR/Copia | 242 | 468658 | 0.81 |
| DNA/DTA | 458 | 346298 | 0.6 |
| DNA/Helitron | 370 | 237796 | 0.41 |
| LTR/unknown | 510 | 207747 | 0.36 |
| DNA/DTC | 116 | 170152 | 0.3 |
| DNA/DTH | 397 | 142366 | 0.25 |
| LTR/Gypsy | 73 | 62300 | 0.11 |
| LINE/CRE | 53 | 32479 | 0.06 |
| MITE/DTH | 103 | 22212 | 0.04 |
| DNA/DTT | 11 | 16527 | 0.03 |
| MITE/DTA | 33 | 9954 | 0.02 |
| MITE/DTM | 26 | 8338 | 0.01 |
| SINE | 18 | 1751 | < 0.01 |
| MITE/DTC | 1 | 507 | < 0.01 |
| Total Genome TEs | 7163 | 5751148 | 9.98 |

**Table 4.** Summary of functionally annotated genes in the *Fusarium oxysporum* strain FO12 genome.

| **Database** | **Annotated Number** | **Percentage (%)** |
| --- | --- | --- |
| Swiss-Prot | 10279 | 64 |
| KEGG | 15643 | 97.4 |
| InterPro | 12693 | 79 |
| Pfam | 10832 | 67.4 |
| GO | 9807 | 61 |
| Annotated | 15667 | 97.5 |
| Total Valid Genes | 16068 | 100 |

The circos plot in Figure 1B highlights the compartmentalised distribution of the FO12 genome, with core chromosomes (Chr01–Chr05 and Chr08-Chr12) harbouring low TE densities, whereas accessory chromosomes (Chr06, Chr07, Chr13, and Chr14) serve as significant reservoirs (Figure 1B-e and 1B-f). In particular, Chr14 represents a hotspot for TEs, with a substantial enrichment of LINE/Tad1, MAGGY, and FoHelitron4, which could drive its structural expansion (Figure 4A). Accessory Chr06 is heavily colonised by Han-full (DNA/DTA) and Tad1 elements, forming a combination distinct from the other accessory compartments. Furthermore, Chr13 is a niche for Fo-Helitron3 and NhORF4-like elements, while the accessory regions of Chr07 are uniquely populated by Fot2, Tfa2, and FoHelitron5 (Figure 4A). These non-overlapping signatures suggest that each accessory chromosome displays a unique history of TE colonisation. Kimura substitution profiling indicates a recent TE proliferation in the accessory chromosomes, consistent with a FO12-specific expansion (Figure 4B). This is further evidenced by a sharp dominance of sequences with extremely low divergence (0–2%), particularly within the LTR/Copia and LINE/Tad1 classes, suggesting an ongoing TE-mediated plasticity of the FO12 accessory genome. In contrast, elements such as Terminal Inverted Repeats (TIR)/CACTA and TIR/Mutator show a higher sequence divergence of around 9%, while for non-TIR/Helitron elements the divergence is 18% to 35%, indicative of lack of recent duplication events for these TEs.

**Figure 4.**
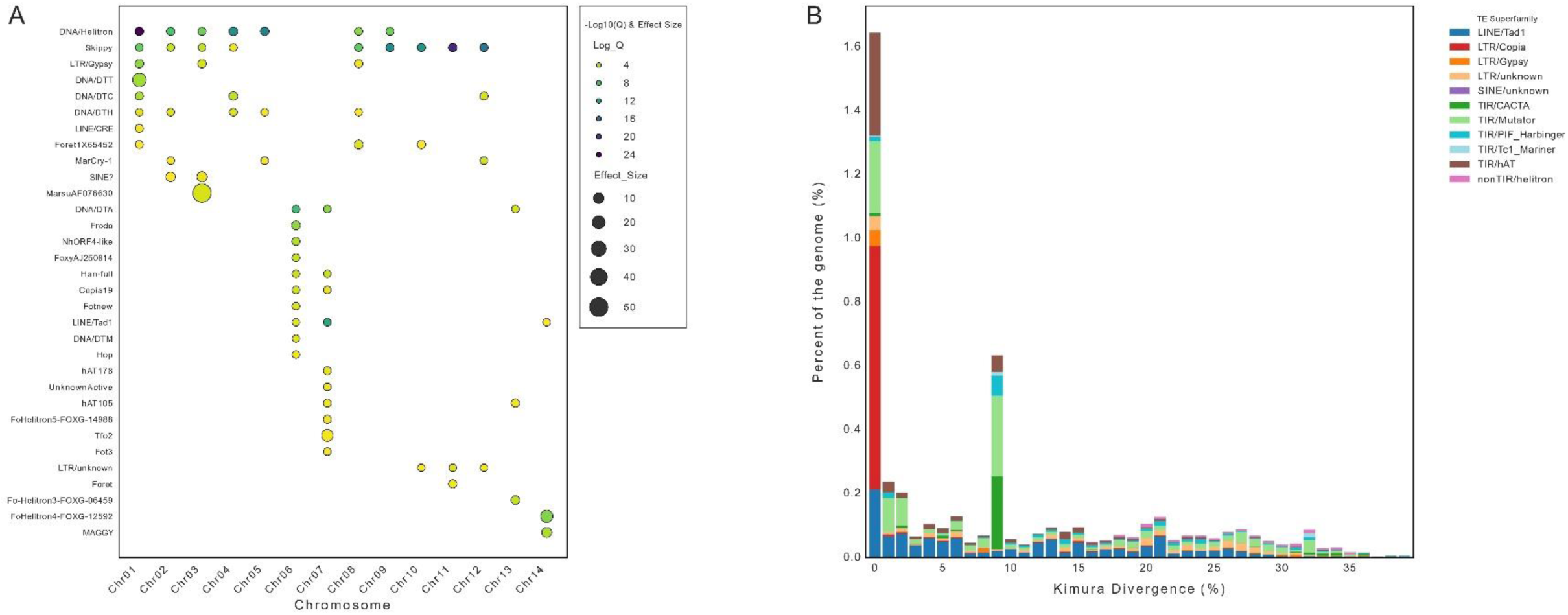
Characterisation of transposable elements across *Fusarium oxysporum* FO12 chromosomes. **(A)** The dot plot illustrates the top 10 enriched families of Transposable Element (TE) for each of the 14 chromosomes the FO12 genome. **(B)** The stacked bar chart shows the distribution of TE sequences according to their Kimura substitution levels (divergence from the consensus sequence). The y-axis shows the percentage of the FO12 genome occupied by each TE class, while the x-axis indicates the Kimura divergence percentage, serving as a proxy for time elapsed since TE duplication events (0% representing very recent events).

Host-specific pathogenicity in *F. oxysporum* has been linked to the presence of lineage-specific virulence effectors such as the Secreted In Xylem (*SIX*) genes^25–32^ We found that, similar to the biocontrol isolate Fo47, the accessory chromosomes of FO12 lack canonical *SIX* genes, with the exception of a putative *SIX8* homolog located on an unpositioned scaffold (Supplementary Table 4-5). The apparent lack of *SIX* genes could explain the absence of virulence of FO12 in every plant host on which this strain has been tested so far. Our analysis using combined results from SignalP, EffectorP and deepTMHMM detected 385 predicted effector candidates encoded in the FO12 genome (Supplementary Table 6). The majority of these putative effector genes (95,85%) are on core chromosomes, with only a smaller fraction (4,15%) located on the accessory genome (Supplementary Image 2). Additionally, comparative genomics showed that this effector collection is highly conserved. Specifically, 241 of the predicted effectors are shared among FO12, the tomato pathogenic reference isolate Fol4287, and the biocontrol strain Fo47 (Supplementary Image 3). However, we also identified a subset of 72 effectors unique to FO12 (Supplementary Image 2-3).

Although FO12 did not cause disease symptoms on any of the plant hosts tested^12,14^, the presence of a large repertoire of putative effectors (Supplementary Table 6-7) is not unexpected, since they may carry out functions unrelated to pathogenicity. Growing evidence suggests that fungal effectors may have originated as antimicrobial proteins^33–35^ that are deployed to manipulate the host microbiota or competing microbes in the soil, as reported in several soil-borne fungi such as *V. dahliae*^36–39^ and *Rosellinia necatrix*^40^. Secondly, core effectors were recently reported in Fol4287 to act as compatibility factors promoting endophytic colonisation of the root cortex^41^. For a soil-dwelling and endophytic organism like *F. oxysporum*, the arsenal of 385 putative secreted effectors may primarily serve to antagonise competing microbes in the rhizosphere and perform asymptomatic colonisation of the plant endosphere. Most of these effector genes are located on the core chromosomes, further supporting the hypothesis of a conserved ecological role (Supplementary Image 3; Supplementary Table 7).

In summary, this study presents the first chromosome-level genome assembly for the biocontrol agent *Fusarium oxysporum* strain FO12, providing a valuable contribution to the genome biology of the *Fusarium oxysporum* species complex. While the *F. oxysporum* species complex has been sequenced extensively, very few genomes to date are supported by proximity labelling data (Hi-C). By providing this kind of assembly, our work offers a high-resolution view of the structural organisation within this lineage. These findings not only advance our understanding of genome compartmentalisation and evolution in *F. oxysporum* but also establish a solid foundation for exploring the molecular mechanisms underlying beneficial plant-endophyte interactions.

## MATERIALS AND METHODS

### Fungal cultivation, DNA extraction, Nanopore library preparation and sequencing

The FO12 strain used in this study was isolated from cork of *Quercus suber* and the original glycerol stock was grown on potato dextrose agar (Difco™ PDA, Becton & Dickinson, USA) medium at 25 °C. For genomic DNA isolation, the strain was then cultivated in 100 mL of homemade potato dextrose broth (PDB) culture in a 500 mL flask for 4 days at 25 °C. Subsequently, mycelium was collected through a centrifugation step at 10000 g for 10 minutes, followed by two wash steps in water and then freeze-dried overnight. High-molecular-weight (HMW) DNA was extracted from the FO12 strain as described by Chavarro-Carrero et al. (2021), except that mycelium was grown in PDB instead of Czapek Dox medium. DNA quality, size, and quantity were assessed using Nanodrop, Qubit, and gel electrophoresis. Library preparation with the Ligation Sequencing Kit (SQK-LSK114) was performed with ∼1.5 µg HMW DNA according to the manufacturer’s instructions (Oxford Nanopore Technologies, Oxford, UK). An R10.4.1 flow cell (Oxford Nanopore Technologies, Oxford, UK) was loaded and run for 24 h, and subsequent base calling was performed using Dorado (version 7.11.2; Oxford Nanopore Technologies, Oxford, UK). To obtain longer reads, ultra-high-molecular-weight (UHMW) DNA, an extra clean-up step was performed. To this end, after DNA precipitation, the Short Reads Eliminator Kit (PacBio, California, USA) was used following the manufacturer’s protocol to select DNA fragments >10 kb.

### *De novo* genome assembly and chromosome-level assembly

To obtain a first draft assembly, passed reads from Nanopore sequencing were used to generate a de novo assembly of FO12 using Flye v.2.9^15^ with -nano-hq -g 50m -m 3000 parameters, to verify the assembly quality and determine the final genomic sequence. Redundant contigs were deleted, and the corrected assembly was generated as output.

To improve the draft assembly, Hi-C sequencing data were generated to verify and produce a chromosome-level genome assembly. Hi-C library construction of *F. oxysporum* strain FO12 was generated according to the standard protocol and sequencing on the Illumina NovaSeq 6000 platform (Phase Genomics Inc., Seattle, WA, USA). Hi-C reads were mapped onto this assembly with Juicer (v2.0) using the “assembly” option to skip the post-processing steps and generate the merged_nodups.txt file^16^. For the juicer pipeline, restriction site maps were generated using the DpnII (GATC) restriction site profile and the assembly was indexed with BWA index (v0.7.17-r1188)^43^ and used to polish the assembly using 3D-DNA^17^. Afterwards, Juicebox (v1.11.08) was used to manually curate the genome assembly. Contigs were merged to scaffolds according to the Hi-C map and Ns were introduced between contigs within scaffolds; gaps between contigs were removed, and contigs were merged.

### Gene annotation and repetitive element analysis

To annotate the *F. oxysporum* FO12 genome, a comprehensive pipeline combining repetitive element masking, *ab initio* gene prediction, and homology-based evidence was employed. First, repetitive sequences and transposable elements (TEs) were *de novo* predicted and classified using the Extensive *de-novo* TE Annotator (EDTA) pipeline v2.2.2^44^. To enhance the accuracy of TE identification, the *de novo* predictions were supplemented with a thoroughly curated TE library derived from the reference strain *F. oxysporum* f. sp. *lycopersici* 4287 (Fol4287)^7^. The resulting consolidated, non-redundant TE library was subsequently used to soft-mask the FO12 genome assembly using the RepeatMasker (Tarailo-Graovac & Chen, 2009) engine integrated into the EDTA framework.

Following repeat masking, structural annotation of protein-coding genes was performed using the Funannotate pipeline v1.8.1^21^. The prediction process integrated *ab initio* gene predictors, primarily utilising AUGUSTUS^46^ and GeneMark-ES^47^, which were trained specifically using *Fusarium* species parameters. To improve structural accuracy, protein sequences from closely related *Fusarium* reference genomes were mapped as homology evidence. The Funannotate pipeline aggregated the *ab initio* and homology-based predictions, generated consensus gene models via EvidenceModeler (EVM)^48^, and robustly filtered out fragmented models or TE-derived artefacts to produce the final comprehensive set of high-quality protein-coding genes.

Protein domains, motifs, and Gene Ontology (GO) terms were assigned by integrating results from InterProScan^22^ and orthology data from eggNOG-mapper v2^23^. Additionally, the pipeline natively predicted carbohydrate-active enzymes (CAZymes) and proteases by querying the dbCAN^49^ and MEROPS databases^50^, respectively, using HMMER3^51^.

### Fungal effector prediction

For effector prediction, EffectorP v3.0^52^ and SignalP v.6.0^53^ were independently used to identify putative secreted proteins from the FO12 predicted proteome, and the overlapping effectors from the two predictions were obtained (Supplementary Table 4-6). The candidate effectorome was then scanned using deepTMHMM v.1.0^54^ to exclude those containing transmembrane helices, yielding the final set of predicted effectors (Supplementary Table 7).

### Data Records

The raw sequencing data of Nanopore and HiC have been deposited in the National Centre for Biotechnology Information (NCBI) under the BioProject number PRJNA1426711 with the accession numbers SRR37327473 and SRR37327474, respectively. The final assembled genome is deposited under the same BioProject at NCBI (GCA_055853645.1), as well as the genome annotations, including CDS and protein-coding regions files.

### Technical Validation

#### Manual correction, validation and evaluation of genome assembly

To obtain a nearly complete and error-free reference genome, we manually corrected the mis-assembly and removed redundant contigs using Hi-C reads alignment within Juicebox visualisation^18^. To remove potential contamination sequences such as mitochondrial genomes and sequencing adaptors, we used the Foreign Contamination Screening (FCS) tool at the time of GenBank submission to remove contaminant sequences to our assembly and contaminant sequences were removed^55^. Furthermore, all centromeres and a few telomeres (4/28) (TTAGGG) are captured via StainedGlass^56^ and trf v.4.09.1^57^ (parameters: 2 7 7 80 10 90 2000 -d -m -l 2).

#### Assessment of assembly quality and completeness

Completeness evaluation of our assembly was performed using BUSCO^58,59^ against the Hypocreales database, yielding a result of 99.6%, indicating a highly accurate and complete assembly. Beyond gene-space evaluations with BUSCO, the base-level accuracy and structural integrity of the assembly were validated using reference-free k-mer spectra analysis via Merqury v.1.3^60^. Comparing the final assembly against a high-accuracy k-mer database (k=21) derived from the Nanopore reads, the FO12 genome achieved an exceptional consensus Quality Value (QV) of 56.52. Furthermore, the overall k-mer completeness was estimated at 99.07%.

Moreover, per-base coverage analysis using independent Oxford Nanopore (ONT) reads demonstrated a uniform depth distribution across both core (mean depth ∼112x, CV < 0.2) and accessory scaffolds (∼110-115x). No significant depth spikes or drops were observed in genomic compartments enriched for LINE/Tad1 transposable elements, confirming the absence of collapsed repeats or misjoins in the accessory genome.

The only substantial depth variation observed (CV > 2.6) was strictly localised to a 28.6 kb terminal block on core Scaffold 4 (positions 1–28,675), where read depth predictably spiked to >2,600x. Sequence homology analysis using BLASTn^61^ identified this region as the collapsed multi-copy ribosomal DNA (rDNA) tandem array, highlighted by alignments to the *F. oxysporum* 18S rRNA gene (e.g., accession LT841236.1). Immediately downstream of this non-coding rDNA locus, structural annotations reveal the onset of canonical protein-coding genes (starting at 29,399 bp), which perfectly coincides with a sharp return to the expected single-copy genomic coverage (∼129x). Together, these multidimensional validation metrics demonstrate that the FO12 assembly is structurally contiguous and that the observed transposable element compartmentalization is a genuine biological feature of its accessory genome.

## Code availability

All commands and pipelines used in data processing were executed according to the manual and protocols of the corresponding bioinformatic software and described in the Methods section, along with the versions. If no detailed parameters were mentioned for the software in this study, default parameters were used as suggested by the developer.

## AUTHOR CONTRIBUTIONS

AD and ALM conceived the project. AD designed the experiments. Phase Genomics Inc. performed HiC library and sequencing. AD and HM analysed the data. ALM and CAB provided the *Fusarium oxysporum* strain FO12 used in this work. CAB, ALM and ADP provided funding. AD, ADP and CAB wrote and revised the manuscript. All authors read and approved the final manuscript.

---

**Supplementary Image 1.**
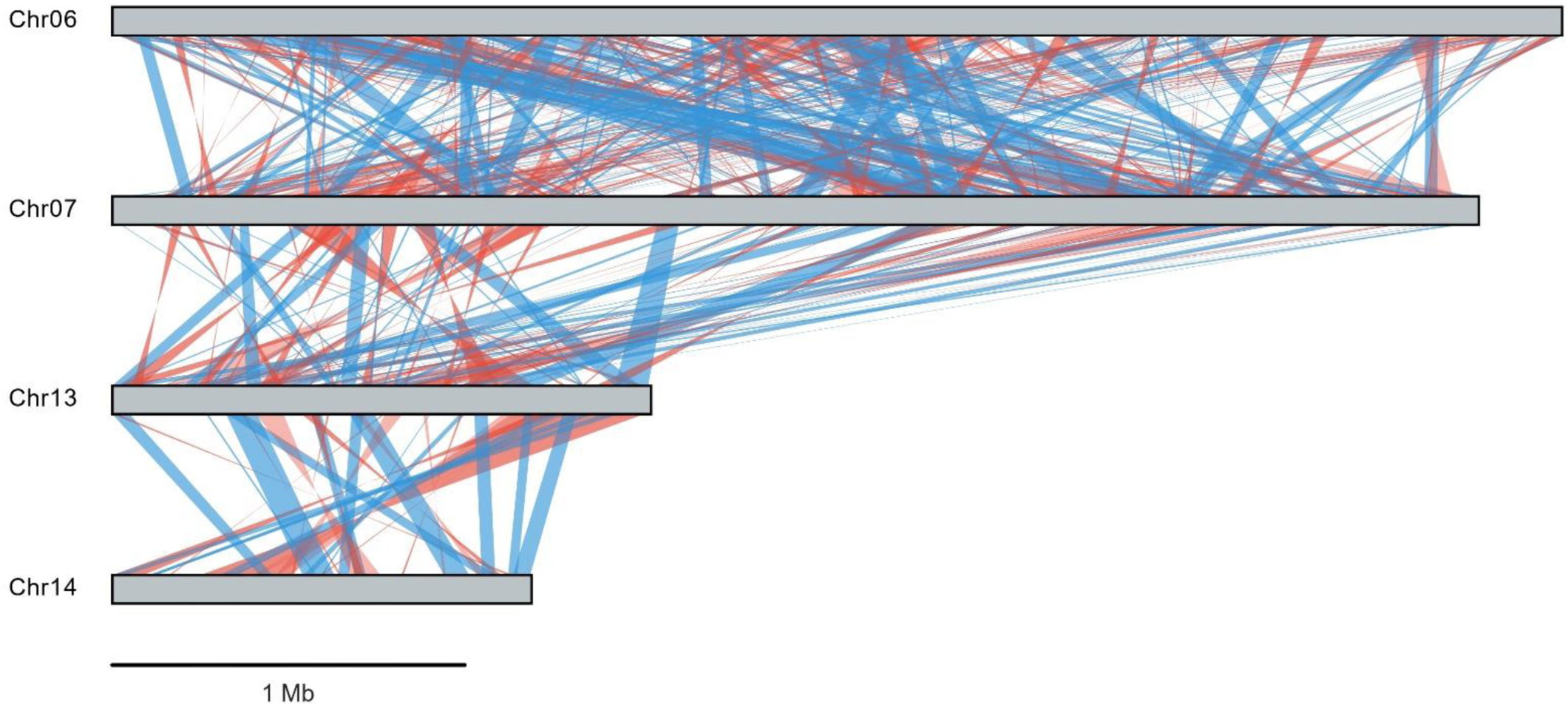
Self-synteny of FO12 accessory chromosomes. Comparative analysis of the four accessory chromosomes (Chr06, Chr07, Chr13, and Chr14) identified in the FO12 genome. Syntenic relationships are represented by colour-coded links; blue lines indicate translocations and red lines indicate genomic inversions.

**Supplementary Image 2.**
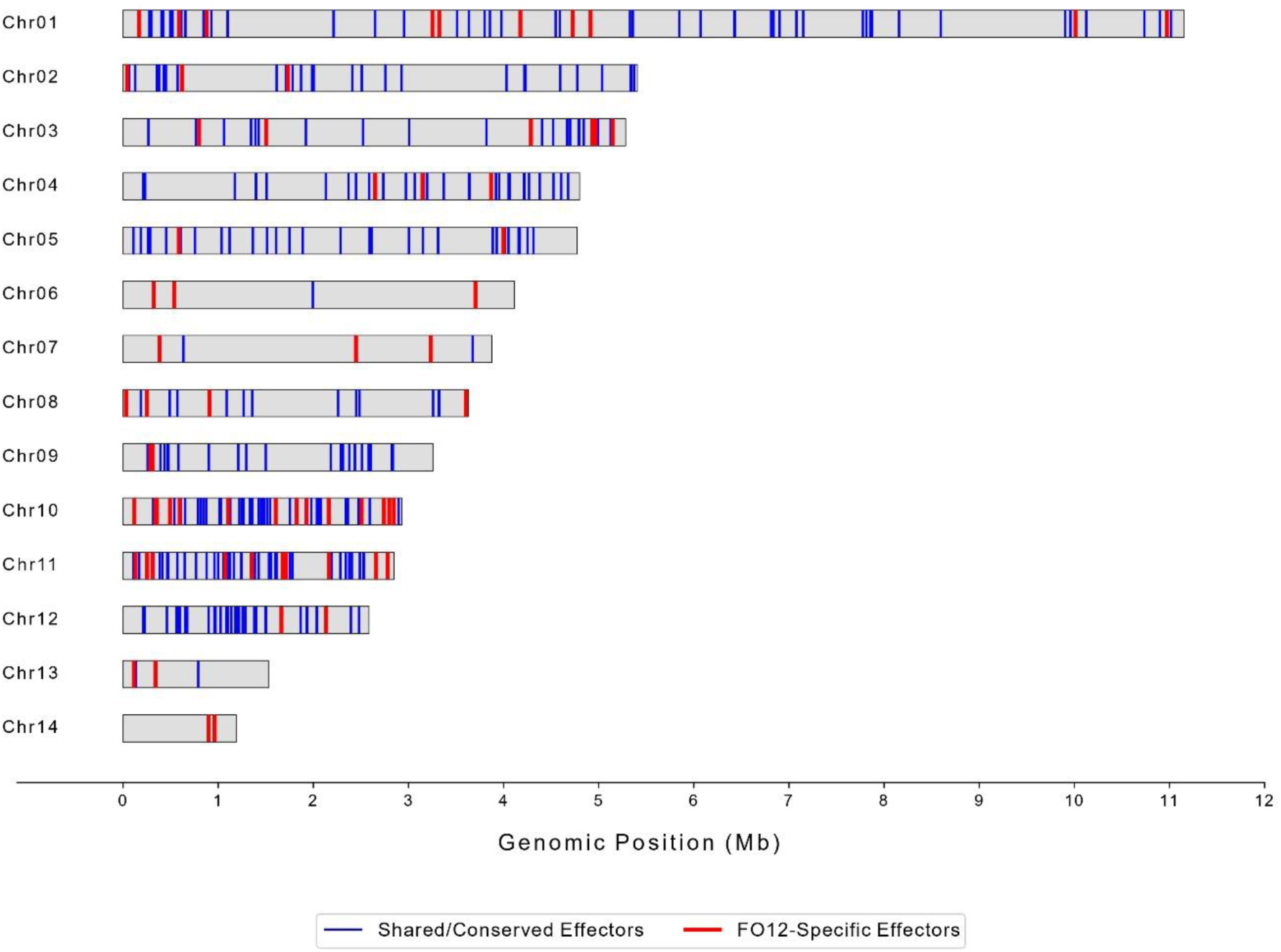
Whole-genome distribution of the FO12 effectorome. Linear genomic map showing the location of the 385 predicted effector genes. Chromosomes are represented by grey bars. Blue vertical lines indicate shared/conserved effectors (found also in Fol4287 and Fo47), while red vertical lines indicate FO12-specific effectors.

**Supplementary Image 3.**
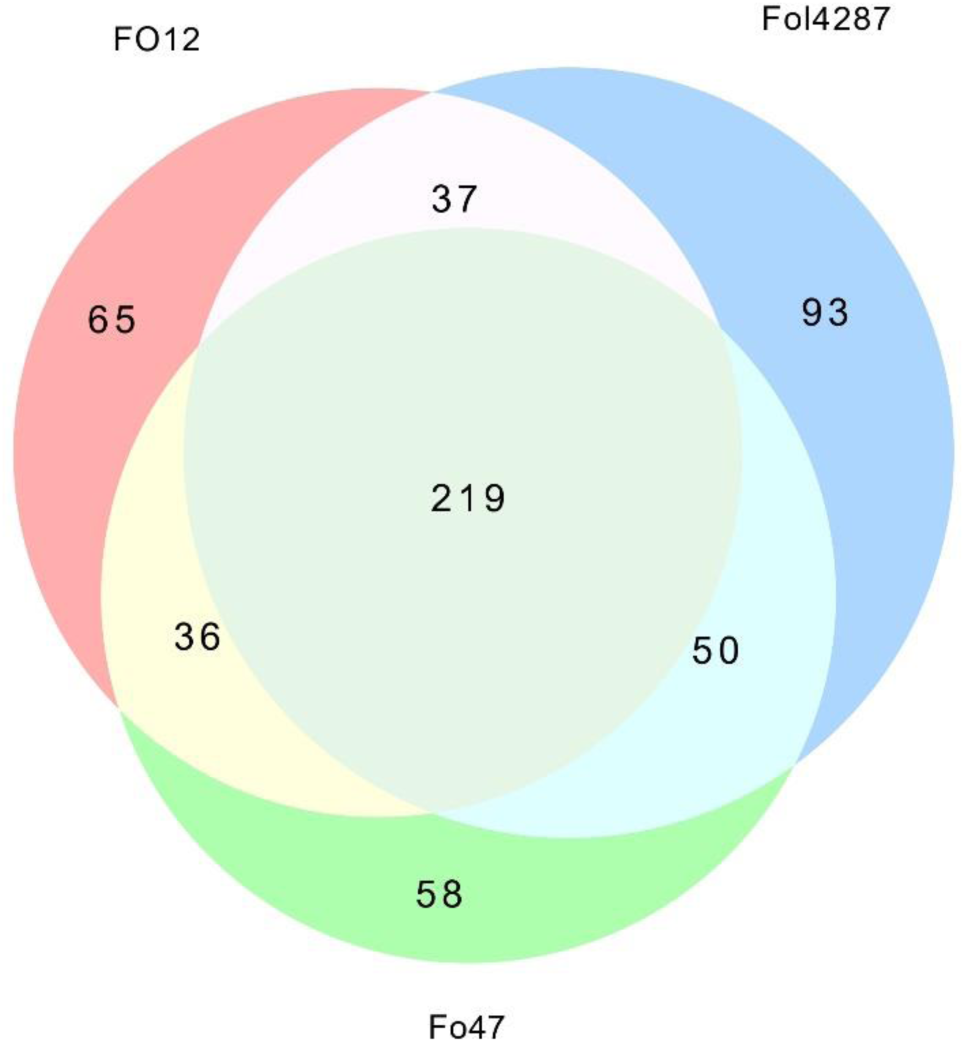
Comparative effectorome analysis and orthology groups. Quantitative summary of the comparative analysis of the predicted effector genes encoded by the FO12, Fol4287 and Fo47 genomes. The numbers represent the clusters of orthologous effector families (identified via CD-HIT) that are either unique or shared between the indicated strains.

